# Dissecting KEAP1-NRF2 interactions: Structural and functional impact of cancer-associated KEAP1 mutants

**DOI:** 10.64898/2026.10.19.760127

**Authors:** Qian Wang, Watanyoo Sopipong, Geoff Wells, Frank Kozielski

## Abstract

Kelch-like ECH-associated protein 1 (KEAP1) is a cysteine-rich substrate adaptor for the CUL3-RBX1 E3 ubiquitin ligase complex, primarily regulating the transcription factor NRF2. By promoting NRF2 degradation under basal conditions, KEAP1 maintains redox homeostasis, while oxidative or electrophilic stress disrupts this regulation, enabling NRF2 activation of cytoprotective genes. Dysregulation of the KEAP1-NRF2 pathway, often through KEAP1 mutations, is implicated in cancer, where constitutive NRF2 activation promotes tumour growth, chemoresistance, and metabolic reprogramming. In this study, we investigated structural and functional consequences of three representative KEAP1 mutants R470C (superbinder), G480W (hypomorphic), and R554Q (null allele) using thermal shift assays (TSA), microscale thermophoresis (MST), isothermal calorimetry (ITC), and X-ray crystallography, complemented by AlphaFold3 modelling. TSA revealed that while inhibitor-induced thermal stability shifts followed similar pattern across mutants, baseline stability varied significantly, with R554Q showing the largest destabilisation. MST and ITC confirmed mutant-specific alterations in NRF2 ETGE peptide binding affinity, consistent with their known functional classifications. Structures of mutant-inhibitor and mutant-peptide complexes showed remarkable structural similarity to native KEAP1, with preserved inhibitor and peptide binding orientations despite residue substitutions. Spatial mapping of reported KEAP1 mutations revealed clustering of superbinder mutations near the IVR interface, while most loss-of-function variants localised within the KELCH domain. Our findings provide a detailed structural and biophysical characterization of clinically relevant KEAP1 mutations, offering mechanistic insights into their functional diversity. This work advances the understanding of KEAP1 mutation effects, with implications for therapeutic targeting of the KEAP1-NRF2 pathway in cancer.

## Introduction

Keap1 is a multifunctional protein that plays a crucial role in cellular defense mechanisms, particularly against oxidative and electrophilic stress to maintain redox homeostasis. It is a cysteine-rich dimeric protein, that consists of 624 amino acids and functions as an adapter for the CUL3-RBX E3 ubiquitin ligase complex. It contains several domains, including the Broad complex Tramtrack and Bric-à-Brac (BTB) domains, the intervening region (IVR), and a C-terminal KELCH domain that is crucial for binding and recognition of its ubiquitination targets (Cuadrado *et al*., 2019; Kopacz *et al*., 2020). KEAP1 is crucial to the post-translational control of nuclear factor erythroid 2-related factor 2 (NRF2), a cytoprotective basic leucine zipper transcription factor that regulates the activity of a battery of genes with antioxidant response elements in their promoter regions. NRF2 regulates the expression of numerous cytoprotective genes, including those involved in glutathione synthesis, thioredoxin-based antioxidant systems, NADH regeneration, and phase I and II detoxification enzymes (Ma, Q., 2013). By controlling cellular NRF2 concentrations, KEAP1 indirectly influences the expression of numerous cytoprotective genes involved in detoxification and antioxidation processes, helping to protect cells from oxidative damage and maintain cellular redox balance.

KEAP1 has the ability to sense oxidative and electrophilic stresses through its 27 cysteine residues, which can be modified by alkylation, complexation or redox reactions (Thang *et al*., 2023; Dinkova-Kostova *et al*., 2002). When exposed to stress, KEAP1 undergoes conformational changes, that inhibit Nrf2 ubiquitination, allowing it to accumulate and translocate to the nucleus where it forms heterodimers with small Maf proteins and activates antioxidant response element (ARE) genes (Baird et al., 2014; Horie et al., 2021). Under normal, unstressed conditions, KEAP1 binds to NRF2 via two motifs in NRF2’s Neh2 domain, a low-affinity DLGex motif and a high-affinity ETGE motif. The interaction between KEAP1 and NRF2 follows a “Hinge-Latch” model, where the ETGE motif acts as a hinge and the DLGex motif as a latch (Horie et al., 2021). Other binding partners of KEAP1 have ExGE-type consensus binding sequences that mimic the tight-binding motif of NRF2. They include sequestosome-1/p62 (pSTGE) and PGAM5 (ESGE) that are involves in autophagy and mitochondrial function respectively, and inhibitor of nuclear factor kappa-B kinase subunit beta (IKKβ) (ETGE) involved in inflammatory and immune signaling responses (Kopacz et al, 2020). These alternative KEAP1 binding partners may be ubiquitinated and degraded, and compete with NRF2 for occupancy of the KELCH domain binding pockets.

Increasing NRF2 transcriptional activity and the expression of ARE genes has been proposed as a therapeutic intervention in a range of conditions spanning inflammation, neurodegeneration and immune function and mitochondrial disorders (Cuadrado *et al*., 2019). Thus, compounds that either mimic oxidative stress (electrophiles that interact with KEAP1 cysteine residues) or interfere with the KEAP1-NRF2 protein-protein interaction have been proposed as therapeutic interventions and a range of compounds exploring both mechanisms have been developed (Crisman et al., 2023). In contrast, in cancer, the KEAP1-NRF2 pathway is frequently dysregulated, resulting in constitutive activation of NRF2 and its downstream effects. In this context sustained high levels of NRF2 results in the expression of ARE-dependent gene products that enable cancer cells to combat oxidative stress and DNA damage induced by chemotherapy (Taguchi, K., Yamamoto, M., 2017). Additionally, NRF2 activation supports the transcription of genes involved in cell growth, proliferation, and apoptosis resistance, thereby promoting tumour growth and survival (Mitsuishi, Y., *et al*., 2012; Scalera, S., *et al*., 2022).

Therefore, KEAP1 as the main regulator of NRF2, is a crucial tumour suppressor protein implicated in cancer development and progression. Loss of heterozygosity (LOH) of KEAP1 is commonly observed in tumours (Teshiba, et al., 2013, Berger et al., 2016). Comprehensive cancer genome studies further highlight the tumour-suppressing role of KEAP1. The Cancer Genome Atlas (TCGA) identified KEAP1 and NRF2 among the most frequently mutated genes in a study of 178 squamous cell lung carcinomas (SQCC), with mutation frequencies of 12% and 19%, respectively. Interestingly, Keap1 and NRF2 mutations are mutually exclusive in SQCC, frequently resulting in a loss-of-function of KEAP1 and unregulated NRF2 activity (Cancer Genome Atlas Research Network, 2012).

NRF2 mutations are focused on the known KEAP1 interaction motifs in the Neh2 domain of the protein, but KEAP1 mutations are distributed throughout its amino acid sequence. Little is known about their functional impact and the functional consequences of these Keap1 mutations are largely unknown. In SQCC, of the 18 observed mutations, two are nonsense mutations (N469fs and P318fs) that truncate the protein, while the remaining 16 ones are missense mutations. Eight of these missense mutations are in the KELCH domain, including R320Q, G333C, D422N, G423V, R470C, G480W, W544C, and R554Q. Western blot analysis of immunopurified Keap1 mutants revealed functional heterogeneity. Mutants such as R554Q, W544C, N469fs, P318fs and G333C lose the ability bind to NRF2, while other retain binding to both NRF2 and CUL3. Notably, a subset of mutant termed “superbinders”, including R320Q, R470C, G423V, D422N, G186R, S243C and V155F, exhibit stronger NRF2 binding than wild-type KEAP1 (Hast, B.E., *et al*., 2014).

In this study, we selected three representative classes of Keap1 mutants for structural and biochemical studies. The R470C mutant was chosen as the representative of a “superbinder”, R554Q was selected as a mutant that fails to bind to NRF2, and G480W was picked as a mutant that binds NRF2 similarly to wild-type KEAP1. We crystallised two of the three Keap1 mutants revealing their structural integrity. We also tested the mutants for their ability to bind to NRF2 peptides and confer resistance to four distinct sulfonamides inhibitors. The results show high structural similarity among mutants and native KEAP1. However, despite structural conservation, G480W and R554Q mutants established relatively lower thermostability compared to the native KEAP1. Furthermore, binding affinity assays also confirm the previous characterisation of mutants. The results also suggest that the decreased thermostability may contribute to diminished ligand-binding capacity of the mutants which might lead to lower efficiency of the inhibitory effect of the inhibitors.

## Materials and Methods

### Subcloning of native Keap1 and selected mutants

The three Keap1 mutants R470C, G480W, and R554Q were generated, based on the previous expression clone described for native Keap1 (Georgakopoulos, N., *et al*., 2022).

### Expression and purification of Keap1 proteins

The expression of native and mutant Keap1 proteins was carried out in E. *coli* BL21-CodonPlus (DE3)-RIPL competent cells (Agilent Technologies, Santa Clara, CA, USA) in Terrific Broth (TB) modified medium (Melford, Chelsworth, UK) supplemented with 50 μg ml^-1^ kanamycin and 34 μg ml^-1^ chloramphenicol. Cell cultures were incubated at 37°C with shaking at 200 rpm until the OD_600_ reached 0.6-1.0. Induction of protein expression was done with 1 mM isopropyl β-D-1-thiogalactopyranoside (IPTG), followed by incubation at 18°C and shaking at 200 rpm for 24 h. Cell pellets were then resuspended in 50 mM HEPES pH 7.5, 300 mM NaCl, 10 mM imidazole, and 1 mM phenylmethylsulfonylfluoride (PMSF) and flash-frozen in liquid nitrogen and stored at −80°C. Cell pellets were thawed at room temperature and lysed by sonication for 30 sec for 10 rounds on ice, with a 60 sec rest period in between. Cell debris was removed by centrifugation at 20,000 rpm for 60 min at 4°C. The supernatant was loaded onto a 5 ml HiTrap TALON crude column (Cytiva, Uppsala, Sweden) equilibrated in 50 mM HEPES pH 7.5, 300 mM NaCl, 10 mM imidazole, and 1 mM PMSF. The column was washed with washing buffer (50 mM HEPES pH 7.5, 300 mM NaCl and 20 mM imidazole), and bound protein was eluted with elution buffer (50 mM HEPES pH 7.5, 300 mM NaCl, and 300 mM imidazole). Sumo-tagged protein was pooled and cleaved overnight with Ulp1 protease by dialysis against 50 mM HEPES pH 7.5, 300 mM NaCl and 1 mM DTT at 4°C. After cleavage, the sumo-tag was removed, and cleaved protein was washed out by washing buffer after the second affinity column (5 ml HisTrap FF Cytiva, Uppsala, Sweden). Samples containing cleaved native Keap1 were pooled and dialysed overnight at 4°C against 50 mM HEPES pH 7.5 and 200 mM NaCl. The protein was then concentrated to 10 mg/ml using an Amicon ultraconcentration device (Millipore), aliquoted, flash-frozen in liquid nitrogen, and stored at −80°C for subsequent use.

### LC-MS to verify the mutants

High-performance liquid chromatography-mass spectrometry was performed to verify the corresponding molecular weight of the proteins. The protein samples were diluted to approximately 1 mg/mL using HPLC grade water prior to HPLC injection for the mass spectrometry. The samples were then passed through a PLRP-S reversed-phase column (Agilent, USA) and eluted with gradient elution mixture of water-acetonitrile. The eluted peaks were subsequently ionised using electrospray ionisation prior to ion separation with a quadrupole time-of-flight tandem mass spectrometry (Agilent, USA). The deconvoluted mass was analysed to determine the protein mass.

### Crystallization of Keap1 in the presence of inhibitors and peptides

Protein crystallisation was conducted on Linbro plates using the hanging drop method. Native Keap1 crystallised at 4°C after 10 days in 3.5 M ammonium acetate and 0.1 M sodium acetate trihydrate pH 4.6. Crystals for mutant R470C appeared in 3.4 M ammonium acetate, 0.1 M sodium acetate trihydrate pH 5.0, and crystals for mutant R554Q appeared in 3.5 M ammonium acetate, 0.1 M sodium acetate trihydrate pH 4.6 after a duration of two weeks. Crystals for the two Keap1 mutants were cryo-protected with 15% ethylene glycol in 1.2-fold of the crystallisation solution and flash-frozen in liquid nitrogen. Two Keap1 mutants were then soaked with four inhibitors (NG268, NG316, NG262 and NG284) at 5 mM for 1 h before the crystals were supplemented with 15% dimethyl sulfoxide (DMSO) in 1.2-fold of the crystallisation solution and flash-frozen in liquid nitrogen. For KEAP1–ETGE peptide complex formation, a 7-residue NRF2 ETGE peptide (DEETGEF) was dissolved in 157 mM ammonium acetate buffer to a stock concentration of 56 mM. Native KEAP1 and its mutants were soaked with the ETGE peptide at final concentrations of 1, 5, and 10 mM for 1 h. Following incubation, crystals were supplemented with 20% DMSO in 1.2-fold of the crystallisation solution and flash-frozen in liquid nitrogen.

### Data collection, processing and refinement

Diffraction data for individual mutant crystals and their complexes with inhibitors and peptides were collected at beamline I03 at Diamond Light Source (DLS). Data were processed with either xia2 3dials or autoPROC to a resolution ranging from 1.7 to 2.4 Å. Structures of the individual mutants R554Q and R470C were determined using molecular replacement (MR) utilizing the published native Keap1 model (PDB entry 3ZGD) via the PHASER-MR project in PHENIX (Liebschner, D., *et al*., 2019). Models were manually optimized using Coot (Emsley, P., *et al*., 2010), and the structures were refined employing PHENIX. With refined mutants R554Q and R470C as models, MR was conducted for the structures of mutants with inhibitors or peptides.

### Structure Prediction of the G480W Mutant Using AlphaFold3

As the G480W mutant could not be crystallised, the amino acid sequence of the human Kelch protein (UniProt primary accession Q14145) was utilised as the reference sequence for structure prediction. The sequence was truncated to a length of 285 residues, corresponding to residues 325 to 605. The glycine residue at position 480 was substituted with tryptophan to generate the mutant sequence. Structure prediction was performed using AlphaFold3 (Abramson J. et al., 2024) via the AlphaFold server (https://alphafoldserver.com/). For reproducibility, the modified sequence was submitted with a constant seed value of 1.

### Thermal shift assays

Thermal shift assays (TSA) were conducted using the Tycho NT.6 instrument (NanoTemper Technologies, Germany) on native and mutated Keap1. This technique was used to determine the inflection temperature of the protein which reflect protein stability in the presence and absence of four inhibitors and peptides. For thermal stability, each mutant and native Keap1 Kelch domain was prepared at 0.5, 1, and 2 mg/mL in final buffer (50 mM HEPES, pH 7.5, 300 mM NaCl). The samples were measured using standard capillaries. After the samples had been loaded into the instrument, they were heated at 30°C/min, from 35°C to 95°C. The protein unfolding was analysed according to the fluorescence intensity ratio at 350 nm and 330 nm (tryptophan/tyrosine shifts). The thermal unfolding inflection temperature (T_i_) was measured in triplicate and the mean values and errors were calculated. The differential scanning fluorimetry was conducted at 0.5 mg/mL protein concentration and a fixed inhibitor concentration of 10 µM. DMSO at 1% concentration was used as a negative control for each protein as the inhibitors were dissolved in DMSO and the stock concentration was 1 mM.

### Microscale thermophoresis

Microscale thermophoresis (MST) was conducted to determine the binding affinities between the Keap1 mutants and each individual inhibitor. The protein was labelled with RED-NHS 2^nd^ Generation dye, which binds to lysine residues on the protein surface. The labelling process was conducted as recommended in the labelling kit manual. After labelling the labelled protein was diluted to 20 nM using MST buffer (50 mM Na_2_PO_4_, 300 mM NaCl, 0.05% Tween-20, and 0.1% PEG8000). The assay was measured at a fixed protein concentration of 10 nM against a 2-fold serial dilution of the ligands, with the highest concentration of 6 µM covering the expected range of the K_d_. The standard capillaries were used to load the samples into the instrument for Binding assays using MO.Control Software and the results were analysed using MO.Affinity Analysis software. Datapoints showing aggregation or adsorption were excluded from the analysis. The results were compared as fold changes relative to that of the native Keap1.

### Isothermal titration calorimetry (ITC)

was performed to determine the binding affinities between the 16-residue ETGE peptide (sequence: AFFAQLQLDEETGEFL) and native KEAP1, including its mutants, using a PEAQ-ITC system (Malvern Panalytical, United Kingdom). The peptide was diluted to a final concentration of 200 µM, and 75 µL was loaded into the syringe. KEAP1, at a concentration of 20 µM, was prepared in a volume of 300 µL and loaded into the sample cell. To eliminate buffer mismatch effects, both the protein and peptide solutions were prepared under the same buffer conditions. Each measurement was conducted in duplicate, and the reported binding affinities represent the average of the two independent experiments.

## Results and Discussion

Three Keap1 mutants and the native protein were expressed and purified for structural and biophysical assays, including TSA, MST, and ITC, to determine both protein thermal stability and binding affinities to inhibitors and NRF2 peptides. Liquid chromatography-quadrupole time-of-flight tandem mass spectrometry (LC-Q-TOF-MS/MS) was employed to determine the deconvoluted mass of the proteins. This was achieved by integrating the chromatographic peak with the corresponding mass spectrum. The deconvoluted mass corroborated well with the calculated molecular weights of each mutant protein, as summarised in Table S1.

### Structure determination of KEAP1 mutants

To investigate structural changes underlying KEAP1 mutations and their interactions with inhibitors and NRF2-derived peptides, both native KEAP1 and its mutants were crystalized and subjected to X-ray diffraction analysis. Well diffracting crystals for mutants R470C and R554Q could be obtained, but unfortunately, the G480W mutant could not be crystallised despite repeated attempts. Native Keap1 has previously been published (Li, X., *et al*., 2004; PDB entry 1U6D) but was also included in the analysis. Therefore, The G480W mutant structure was predicted using AlphaFold3. The model yielded an overall per-atom confidence estimate score (pLDDT) above 90 and the predicted template modeling score (pTM) of 0.96 which are considered indicative of a highly reliable and high-quality prediction. The overall conformation of the predicted G480W structure closely resembles the X-ray crystallographic structures of native KEAP1 and other mutants, suggesting no major deviations in fold. Crystallographic statistics for experimentally determined native and mutant structures are summarised in Table S2.

### Comparison reveals high structural similarity among mutants and native KEAP1

The KELCH domain structure of native KEAP1 (PDB entry 1U6D), is composed of six distinct β-propeller subdomains (Fig 1A). The experimentally determined structures of R470C and R554Q, together with the AlphaFold3-predicted model of G480W, all exhibit a very high degree of structural similarity to native KEAP1 as shown in Figs 1B-D. Hydrogen bond analysis, however, revealed subtle local changes at the mutation sites. In the R470C mutant, the substitution of Arg with Cys abolished key hydrogen bonds between the guanidium group of Arg and the carboxyl groups of Asp422 and Glu493. In the mutant, the cysteine’s sulfhydryl group instead engages in hydrogen bonding with nearby water molecules (Fig 1E). Since this residue is located on the basal surface of the KELCH domain—opposite the ligand-binding site—this mutation may confer increased local flexibility, potentially facilitating ligand accommodation or allosteric effects. The G480W mutant showed no major conformational changes at the mutation site, as visualised in the AlphaFold3 model (Fig 1F). Although tryptophan is bulkier than glycine and typically introduces structural rigidity, the predicted model maintains the loop conformation at position 480. However, as this model is computationally predicted and not experimentally validated, further crystallographic evidence is required to confirm this structural interpretation. In contrast, the R554Q mutation led to a more pronounced disruption of local interactions. The loss of the arginine guanidinium group abolished hydrogen bonds with the hydroxyl group of Ser580, the carboxyl group of Glu582, and the backbone atoms of Gly570 and Ser580 (Fig 1G). These disruptions may destabilise the local architecture and affect the protein’s ability to bind NRF2 or other partners.

**Figure 1.**
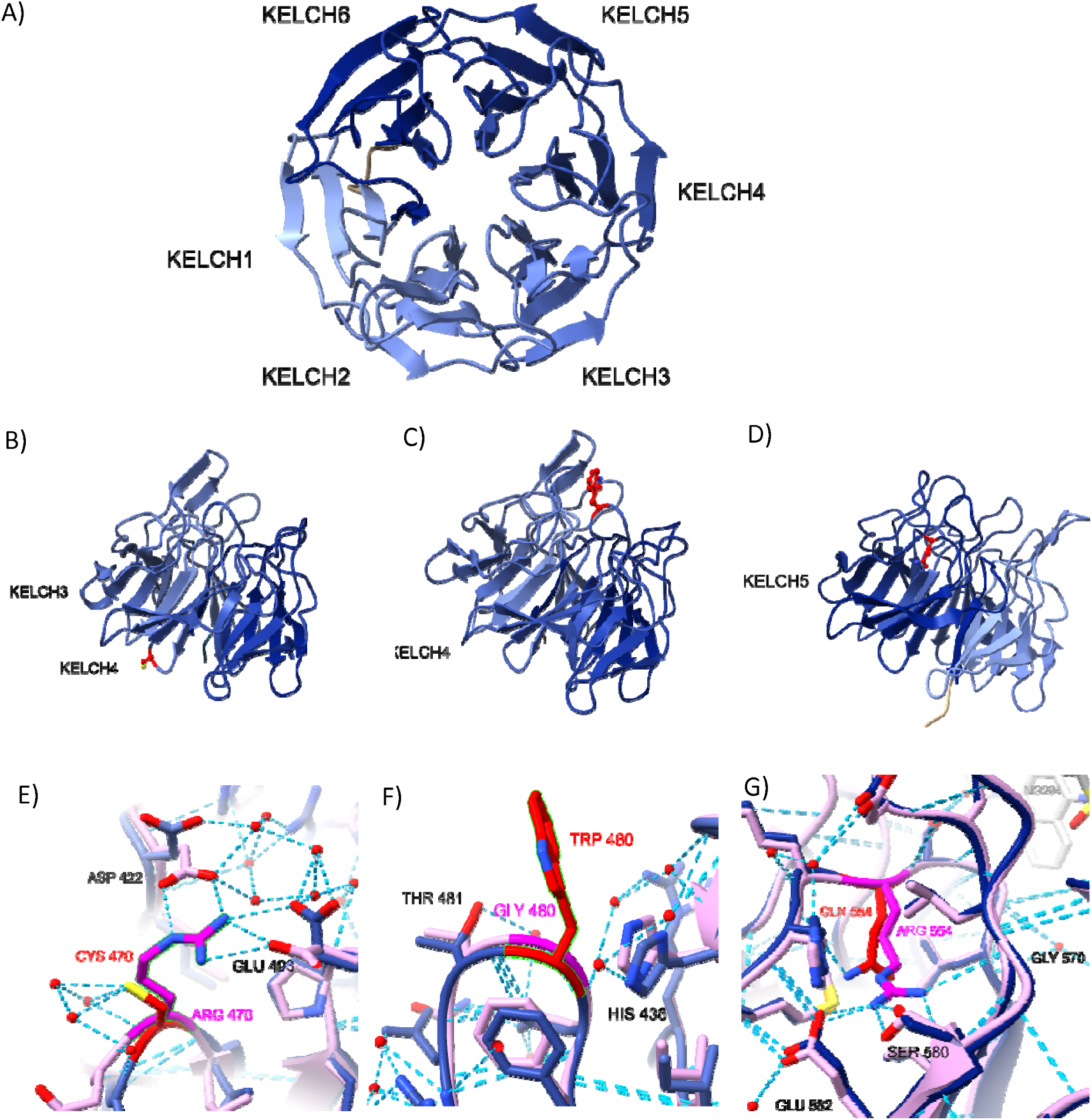
Keap1 KELCH domain structure. **A)** Native Keap1 (PDB entry 1U6D) KELCH 1 (aa 327-372), KELCH 2 (aa 373-423), KELCH 3 (aa 424-470), KELCH 4 (aa 471-517), KELCH 5 (aa 518-564), KELCH 6 (aa 565-611); **B)** Keap1-R470C mutant. **C)** Keap1-G480W mutant. **D)** Keap1-R554Q mutant. Mutant residues are shown as ball-and-stick model and highlighted in red colour. The bottom panel is a close-up view of mutant residues and their interactions with neighbouring residues; **E)** R470C, **F)** G480W, **G)** R554Q. The mutant residues and native residues are coloured in red and magenta, respectively. Hydrogen bonds are represented by blue dashed lines.

### Reduced Thermal Stability in Keap1 Mutants

The thermal shift assays revealed significant differences in thermal stability across Keap1 mutants, compared to native Keap1. Inflection temperatures (T_i_), determined by the fluorescence intensity ratio at 350 nm/330 nm, are presented in Fig 6A and summarised in Table 1. Native Keap1 exhibited a T_i_ of 64.9 ± 0.05°C and the R470C mutant displayed a thermal stability very similar to that of native Keap1. In contrast, the G480W mutant exhibited a moderate reduction in T_i,_ with a 5°C decrease. The R554Q mutant was the least thermally stable, showing a substantial T_i_ reduction of 15.5°C relative to native Keap1. The pronounced reduction in thermal stability observed in the R554Q mutant may be explained by structural disruptions at the interface of Kelch subdomains. R554 is located near the boundary of the fifth and the sixth Kelch subdomain, where its guanidium side chain forms hydrogen bonds with several key residues of Ser580, Glu582 and the Gly570 backbone (Figs 1D,G). The substitution to glutamine eliminates these interactions, potentially weakening the inter-blade packing and destabilizing the local β-propeller architecture. This destabilization may propagate across the domain, contributing to the significantly lower T_i_ observed. In contrast, the G480W mutation induces more subtle local perturbations without disrupting subdomain interactions, resulting in only a moderate reduction in thermal stability. In summary, although the structural integrity is unchanged in the mutant structures, there are significant differences in the thermodynamic stability of G480W and R554Q mutants.

**Table 1.**
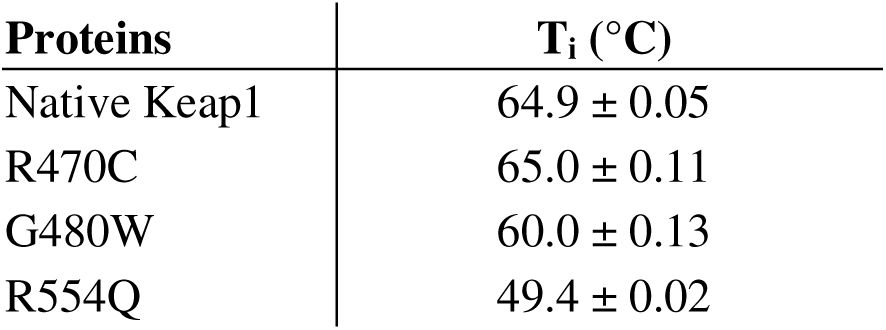
Inflection temperature of native Keap1 and its mutants.

| Proteins | $T_i$ ($^\circ\text{C}$) |
| --- | --- |
| Native Keap1 | $64.9 \pm 0.05$ |
| R470C | $65.0 \pm 0.11$ |
| G480W | $60.0 \pm 0.13$ |
| R554Q | $49.4 \pm 0.02$ |

### KEAP1–ETGE peptide complexes show conserved binding geometry across native and mutant Keap1

A similar structural pattern was observed for the KEAP1–ETGE peptide complexes. Crystals of native KEAP1, along with R470C and R554Q mutants, were soaked with a seven-residue ETGE peptide (DEETGEF) to investigate the potential impact of mutations on peptide recognition. Overlay of each mutant–peptide complex with the native KEAP1–peptide complex revealed highly conserved binding conformations (Fig 2). Despite the point mutations, no significant deviations in the peptide binding orientation or overall domain architecture were observed. The ETGE motif adopted nearly identical conformations in all complexes, and the KELCH domain fold remained structurally intact. These results suggest that the observed differences in binding affinity among mutants are not due to altered peptide accommodation or binding-site disruption, but likely stem from subtle changes in interaction networks or local flexibility.

**Figure 2.**
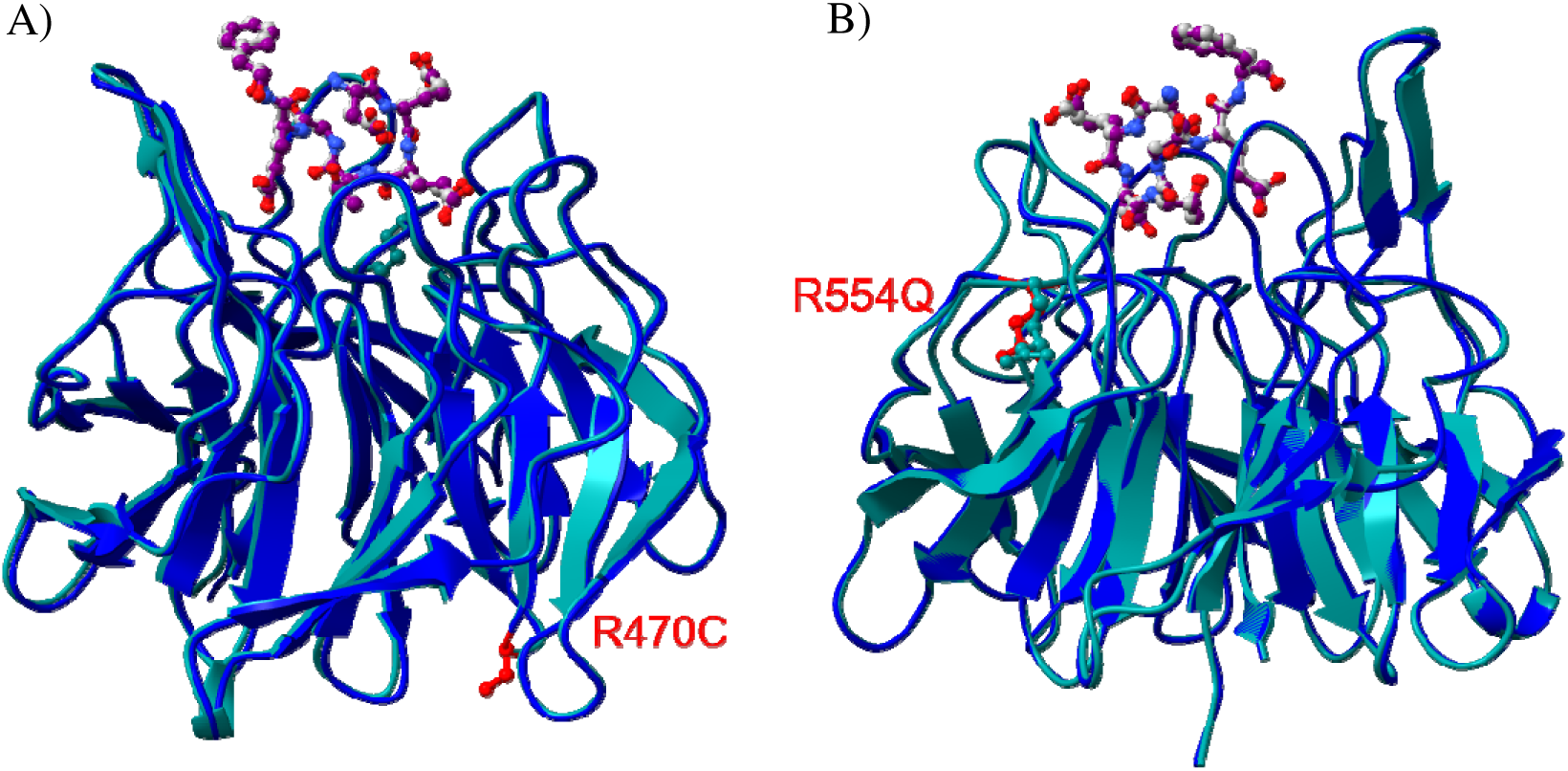
Superimposition of mutant KEAP1–ETGE peptide complexes with native KEAP1-peptide complexes reveals conserved peptide binding. Native KEAP1 is shown in dark cyan and mutants are shown in medium blue. The ETGE peptide (residues DEETGEF) is displayed as a ball-and-stick representation. The ETGE peptide is coloured in grey and purple in native and mutant KEAP1, respectively. A) R470C. B) R554Q.

### Binding affinity analysis confirms mutation characteristics

Using MST, a 16-residue NRF2 peptide containing the ETGE motif was titrated as a ligand against fluorescently labelled native KEAP1 and its mutants (Fig 3A). The NRF2 ETGE peptide bound to native KEAP1 with a K_d_ of 170.8 ± 48.7 nM (Fig 3C). This value is approximately sevenfold higher than previously reported K_d_ of 23.9 nM for the same ETGE peptide measured using surface plasmon resonance (SPR) (Abed, D.A., *et al*. 2015). The R470C mutant exhibited enhanced binding affinity, with a K_d_ of 77.4 ± 14.9 nM, indicating a stronger interaction than that of the native protein. The G480W mutant showed comparable binding affinity to native KEAP1, with a K_d_ of 189.7 ± 102.0 nM. In contrast, the R554Q mutant displayed a substantial loss of binding affinity, with a K_d_ of 669.5 ± 198.6 nM, representing an approximately fourfold reduction in affinity relative to the native protein. These MST-derived binding profiles are consistent with the established functional classifications of the mutants: R470C as a superbinder, R554Q as a null allele, and G480W as a hypomorphic mutant, each exhibiting distinct NRF2 interaction profiles.

**Figure 3.**
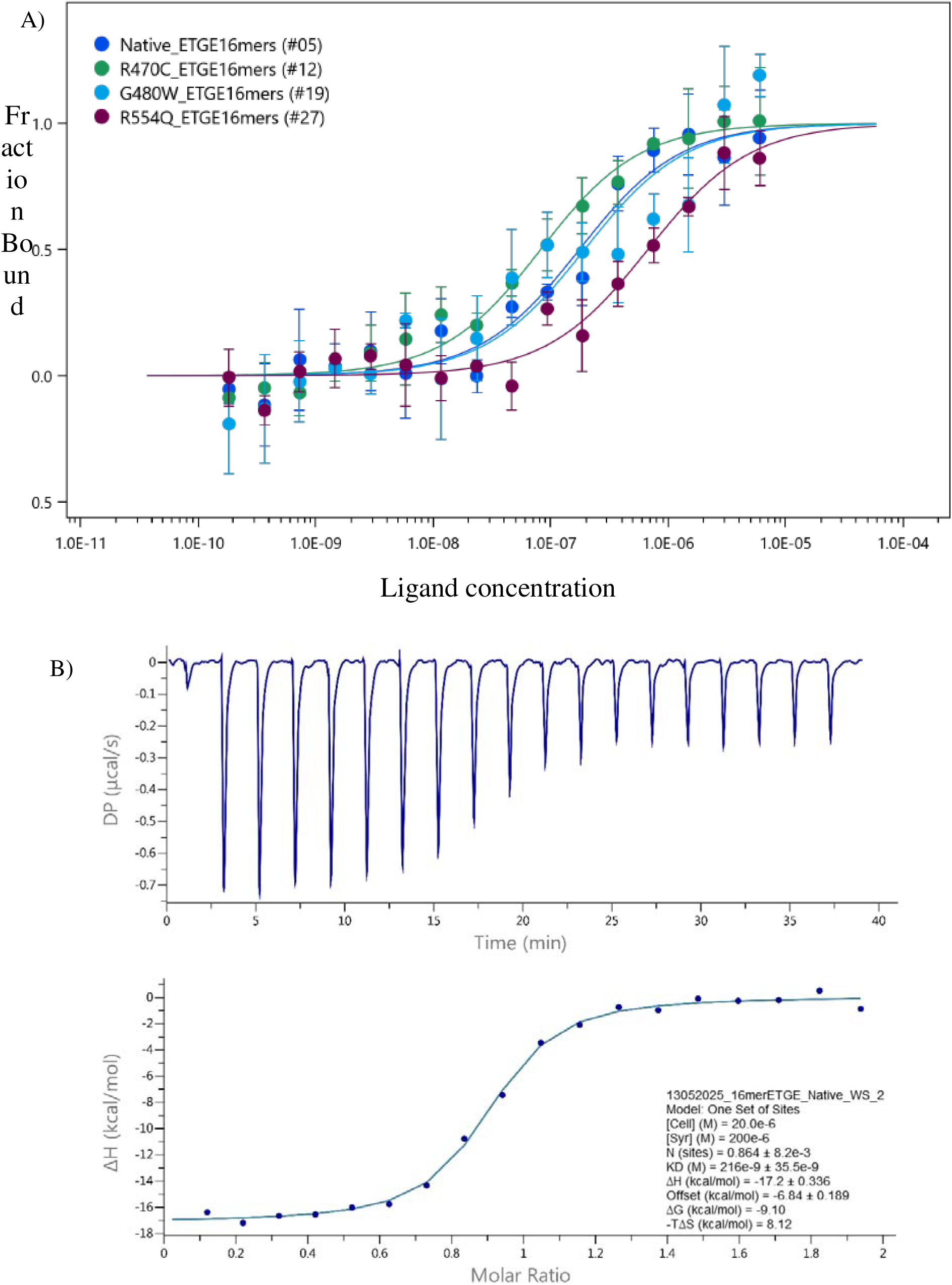
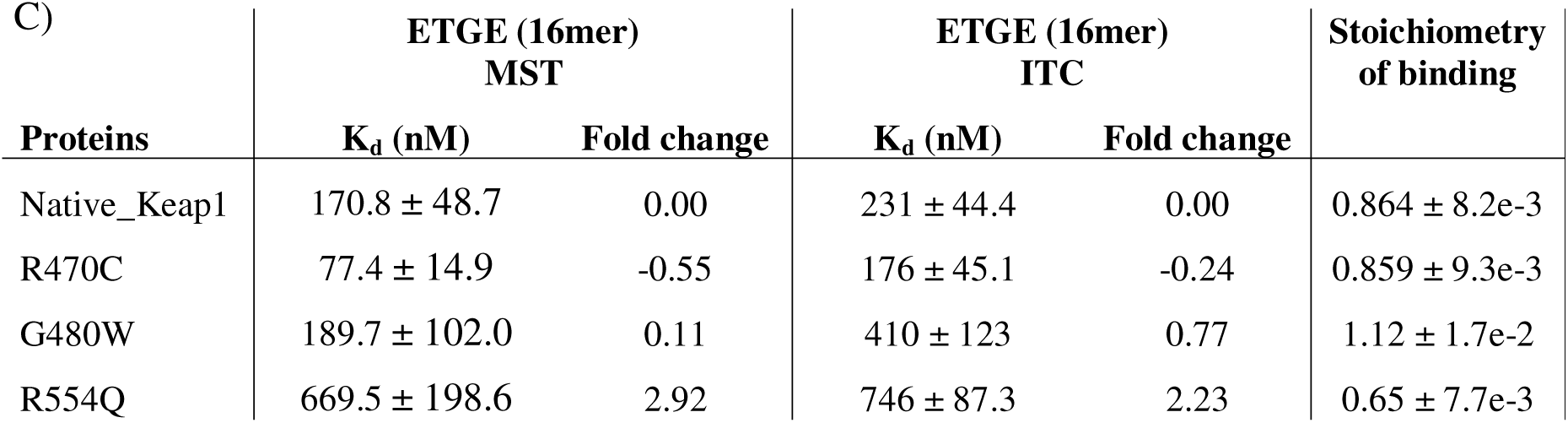
A) MST dose-response curves of native Keap1and its mutants with the 16-residue ETGE peptide. B) ITC titration curves of native KEAP1 with the 16-residue ETGE peptide. C) Summary of binding affinities of the 16mer NRF2 ETGE peptide for native and mutant Keap1 proteins.

To further validate these findings, ITC was performed using the same 16-residue ETGE peptide. The results reflected similar trends to the MST analysis, although the absolute K_d_ values were generally higher, as is typical for this method (Winiewska, M., *el al.*, 2017). Each measurement was conducted in duplicate, and average values are reported (Fig 3B-C). Native KEAP1 exhibited a K_d_ of 231 ± 44.4 nM, while the R470C mutant again showed enhanced affinity, with a K_d_ of 176 ± 45.1 nM. The G480W mutant displayed a weaker interaction than observed by MST, with a K_d_ of 410 ± 123 nM, nearly a two-fold decrease. The R554Q mutant demonstrated a consistent loss of binding affinity across both assays, with a K_d_ of 746 ± 87.3 nM. Despite minor differences in absolute values, the ITC results corroborate the MST data, reinforcing the distinct binding behaviour of the three KEAP1 mutants.

### KEAP1–inhibitor complexes reveal conserved binding orientations across mutants

To investigate the impact of mutations on small-molecule binding, crystal structures of Keap1 R470C and R554Q mutants were determined in complex with four sulfonamide inhibitors NG262, NG268, NG284, and NG316 (Fig 4) and compared with the corresponding native KEAP1–inhibitor complexes. Overlay analyses of the mutant structures with native KEAP1 revealed that the overall fold of the KELCH domain remained highly conserved, and the inhibitors occupied nearly identical binding pockets across all structures (Figs 5–6). Despite local residue substitutions, all four inhibitors adopted remarkably similar binding orientations in the mutant proteins compared to the native structure. This suggests that the amino acid substitutions at positions 470 and 554 do not induce substantial rearrangement of the ligand-binding site or disrupt key protein–ligand interactions. These findings support the hypothesis that the functional effects of these mutations may arise from subtler factors, such as dynamic flexibility or altered affinity, rather than gross structural perturbation of the binding pocket.

**Figure 4.**
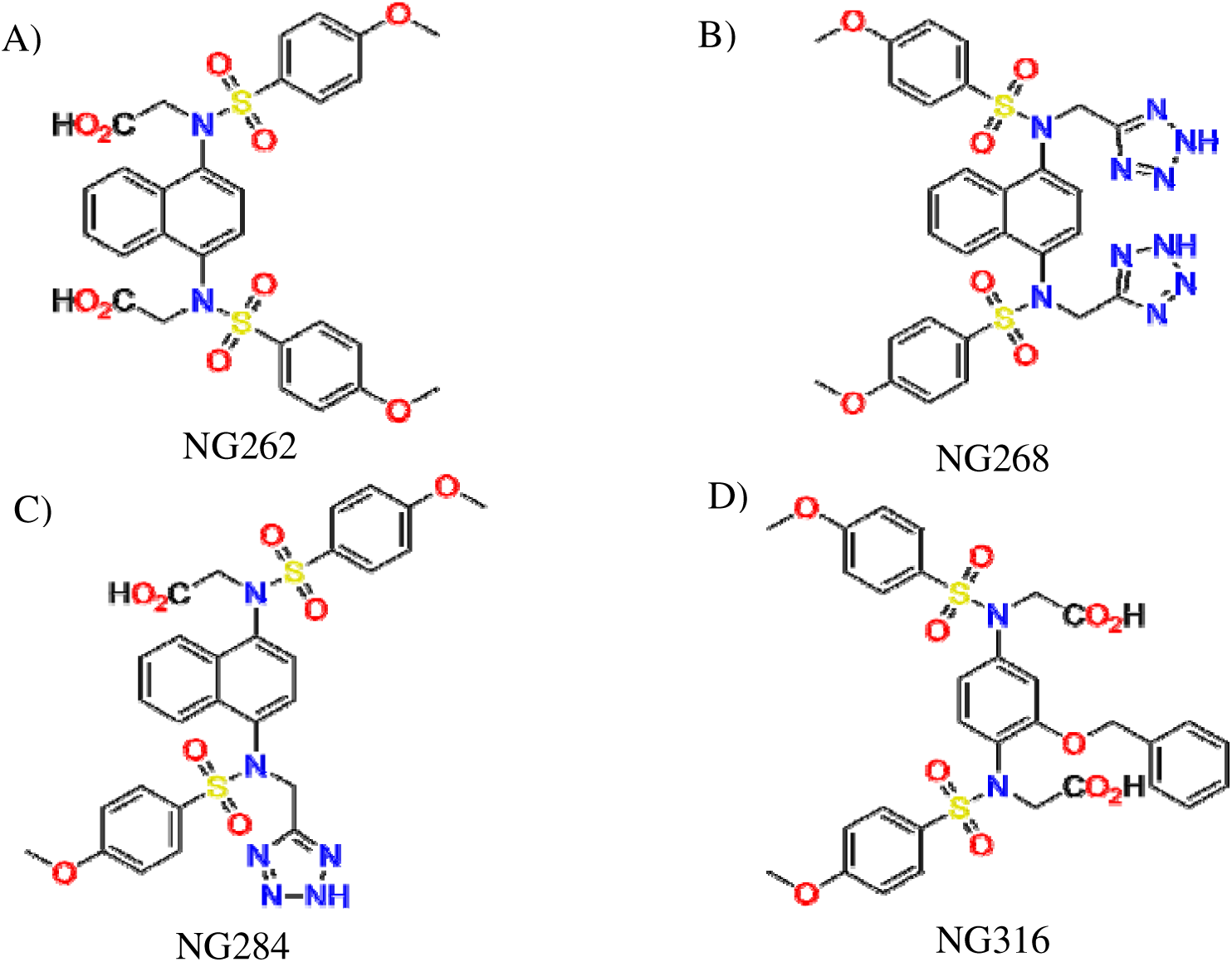
The chemical structures of the four inhibitors A) NG262; B) NG268; C) NG284; D) NG316.

**Figure 5.**
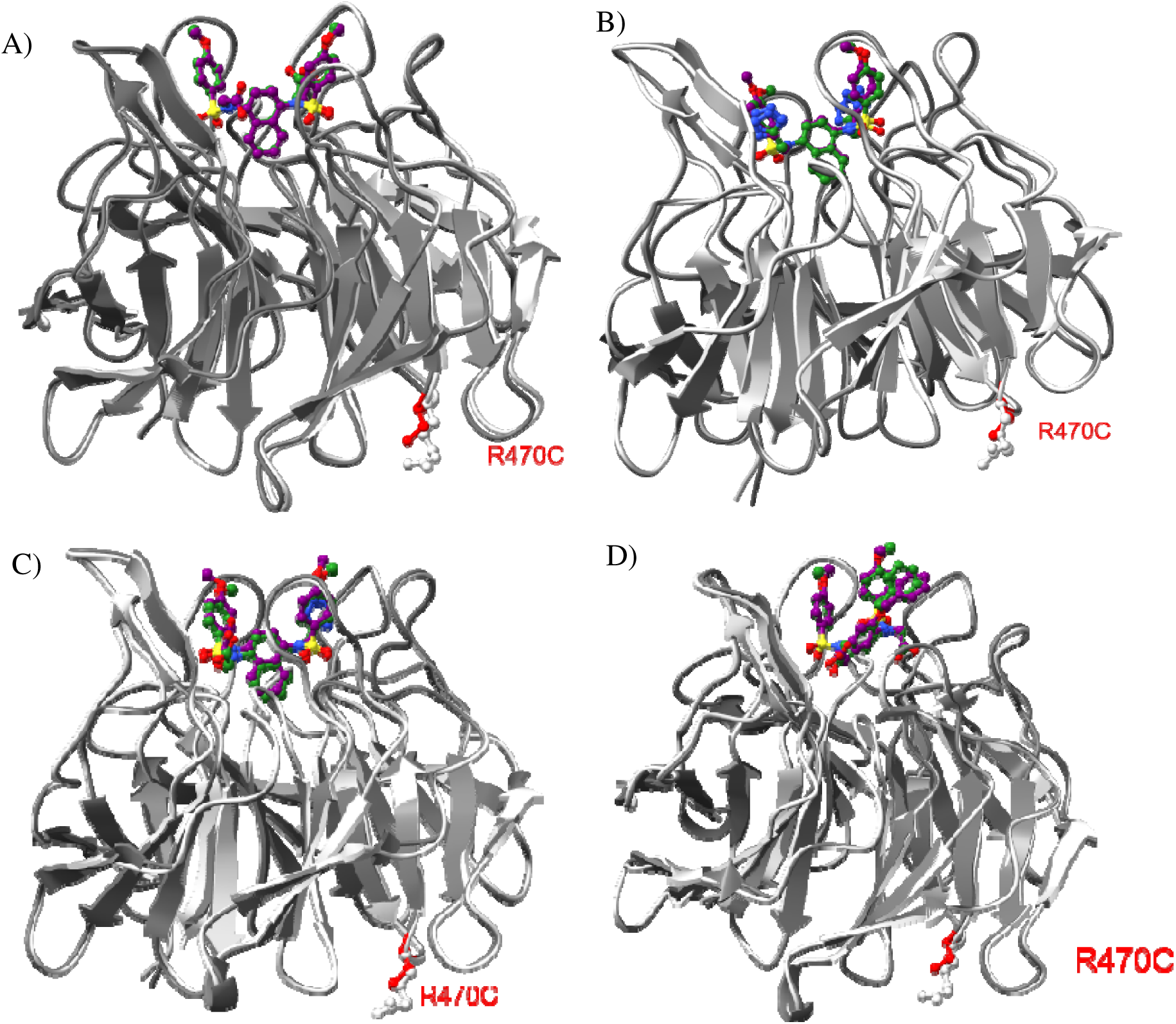
Overlay of Keap1 KELCH domain-inhibitors complexes. Each panel displays the structural comparison between native KEAP1 (light grey) and R470C mutant protein (shown in dark grey) in complex with four inhibitors. Mutated residues are highlighted in red. The inhibitors in native KEAP1 model and mutant model are coloured in green and purple respectively; A) NG262; B) NG268; C) NG284; D) NG316.

**Figure 6.**
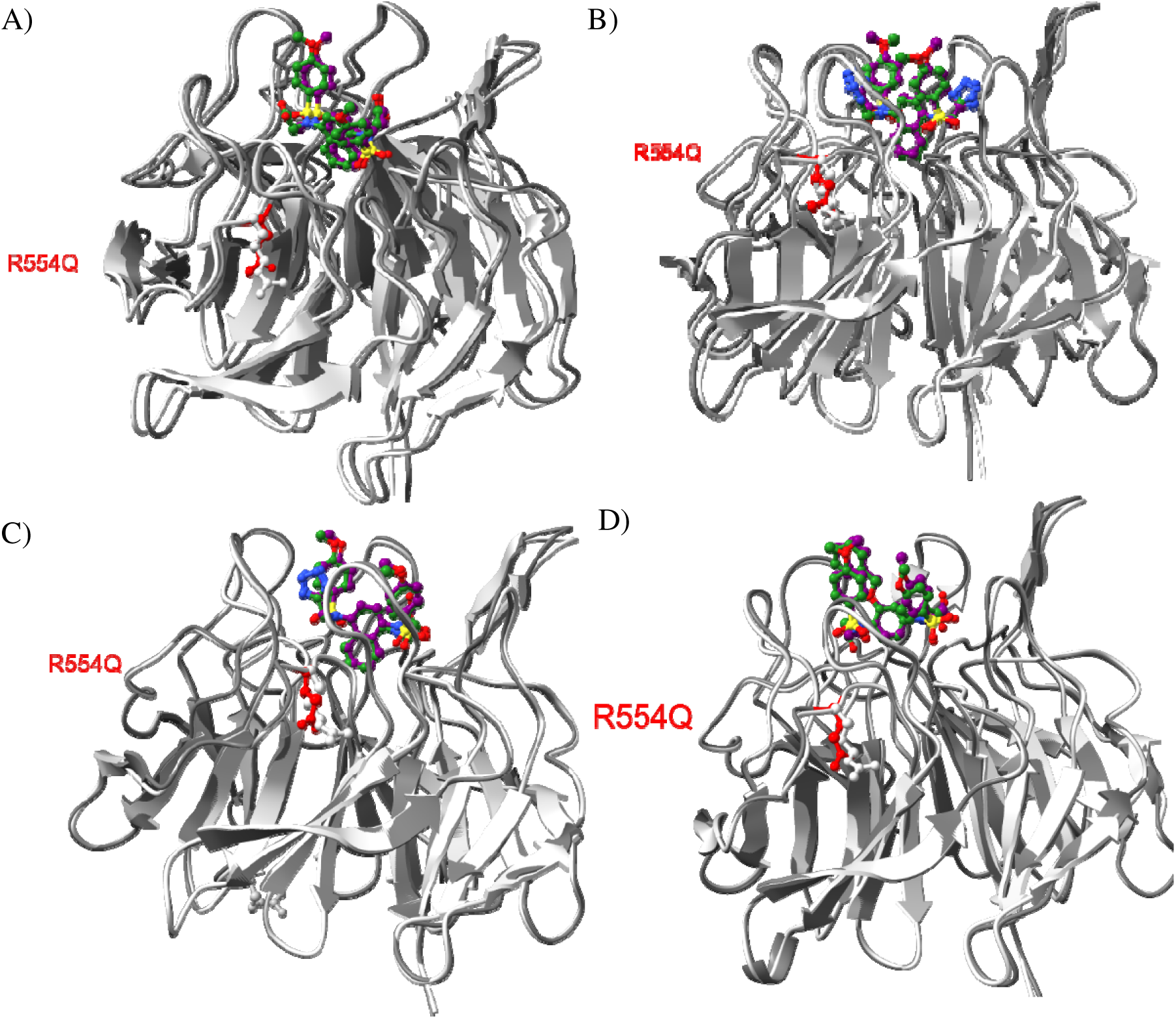
Overlay of Keap1 KELCH domain-inhibitors complexes. Each panel displays the structural comparison between native KEAP1 (light grey) and R554Q mutant protein (shown in dark grey) in complex with four inhibitors. Mutated residues are highlighted in red. The inhibitors in native KEAP1 model and mutant model are coloured in green and purple respectively; A) NG262; B) NG268; C) NG284; D) NG316.

### KEAP1 mutants exhibit differential baseline stability but preserve inhibitor response profiles

We first employed TSA to investigate the impact of mutants on inhibitor binding (Fig 7). NG284 induced the largest increase in T_i_ for native Keap1, raising it by 16.1°C, followed by NG262, which increased the T_i_ by 12.9°C. NG316 caused a moderate shift of 2.8°C, while NG268 only slightly lowered the T_i_ by 0.3°C, relative to the control (1% DMSO). Notably, the R470C mutant exhibited a T_i_ range comparable to that of native Keap1. In this mutant, NG284 and NG262 increased T_i_ by 16.6°C and 13.4°C, respectively, while NG316 shifted the T_i_ by 2.9 °C. NG268 did not significantly alter T_i_ compared to the control. The G480W mutant showed generally lower T_i_ values than R470C, with a baseline T_i_ approximately 5°C lower than native Keap1. NG284 elevated the T_i_ by 15.7°C, and NG262 by 12.9°C. NG316 exerted a notable influence on G480W, raising the T_i_ by 4.2°C, while NG268 caused a slight T_i_ decrease of 0.2°C. In contrast, the R554Q mutant exhibited the most substantial reduction in T_i_ relative to native Keap1, with a 15.7°C decrease in the control. NG284 had the most pronounced impact on R554Q, increasing the T_i_ by 17.6°C, followed by NG262 and NG316, which increased the T_i_ by 12.4°C and 2.4°C, respectively. NG268 did not alter the T_i_ of R554Q. Overall, these findings indicate that NG284 consistently induced the largest T_i_ shifts across all proteins, while the extent of T_i_ modulation varied significantly depending on the specific mutant and inhibitor combination. In summary, inhibitor-induced shifts in T_i_ followed a consistent pattern across all mutants, irrespective of specific residue alterations. Although each mutant had a unique T_i_ compared to native Keap1, the overall T_i_ shift pattern remained similar.

**Figure 7.**
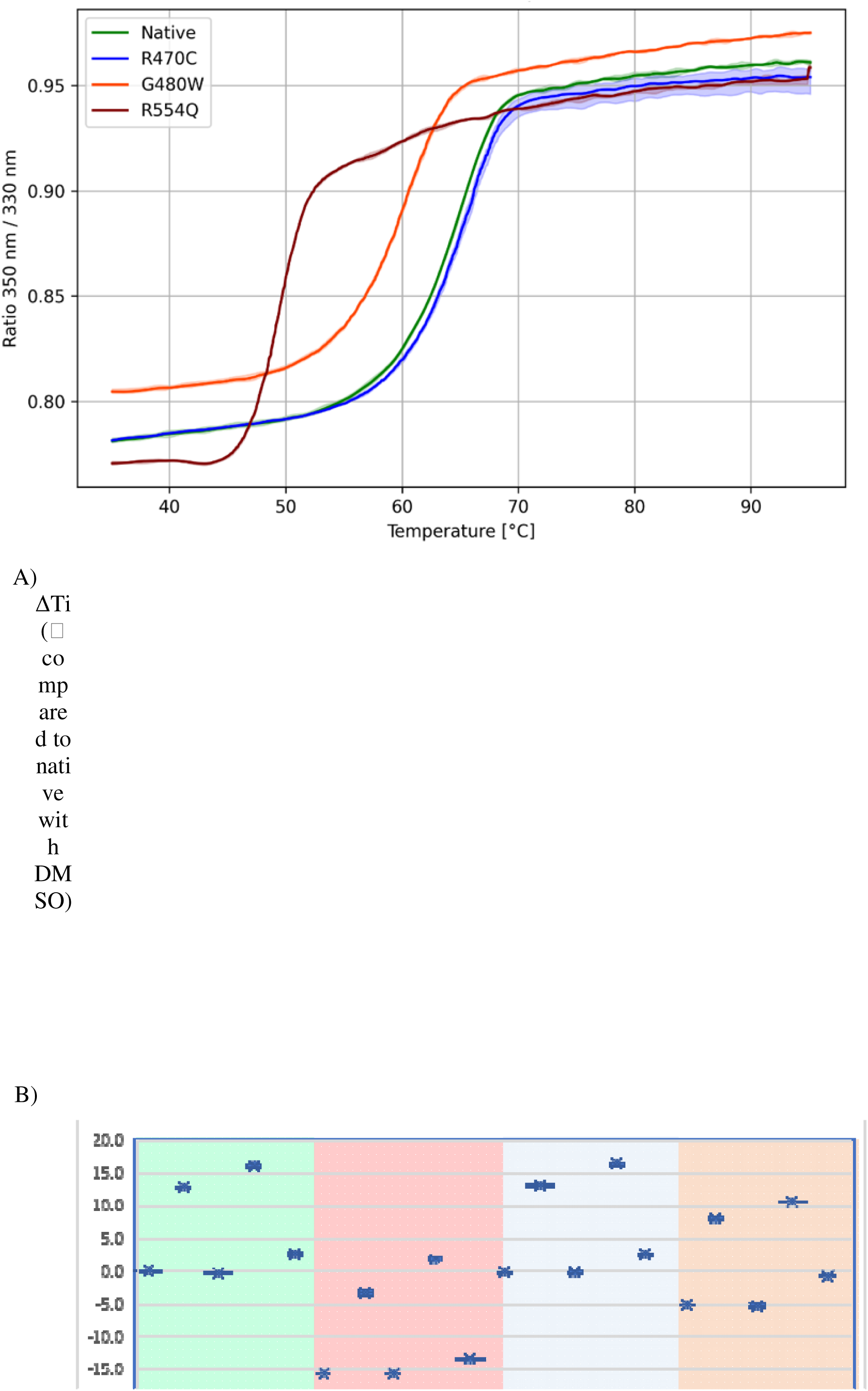
A) The fluorescence intensity ratio 350 nm / 330 nm profile comparison. The lines corresponding to the average 350 nm / 330 nm from the triplicate, while the shaded region represents min-max values from the triplicate. B) Inflection temperature comparison of mutants to native Keap1 in the presence of DMSO.

### Binding affinity analysis reveals differential inhibitor affinities among mutants

MST was employed to measure binding affinities, revealing altered inhibitor affinities for the G480W and R554Q mutants. Ligand-dependent changes were plotted by displaying Fnorm values as a function of ligand concentration in a dose-response curve (Fig 8). Fnorm was calculated as the ratio of F1 (fluorescence in the heated state) to F0 (fluorescence in the cold state, pre-IR laser activation). Binding affinity measurements were conducted at the shortest MST-Ontime (1.5 sec) (Fig 8).

**Figure 8.**
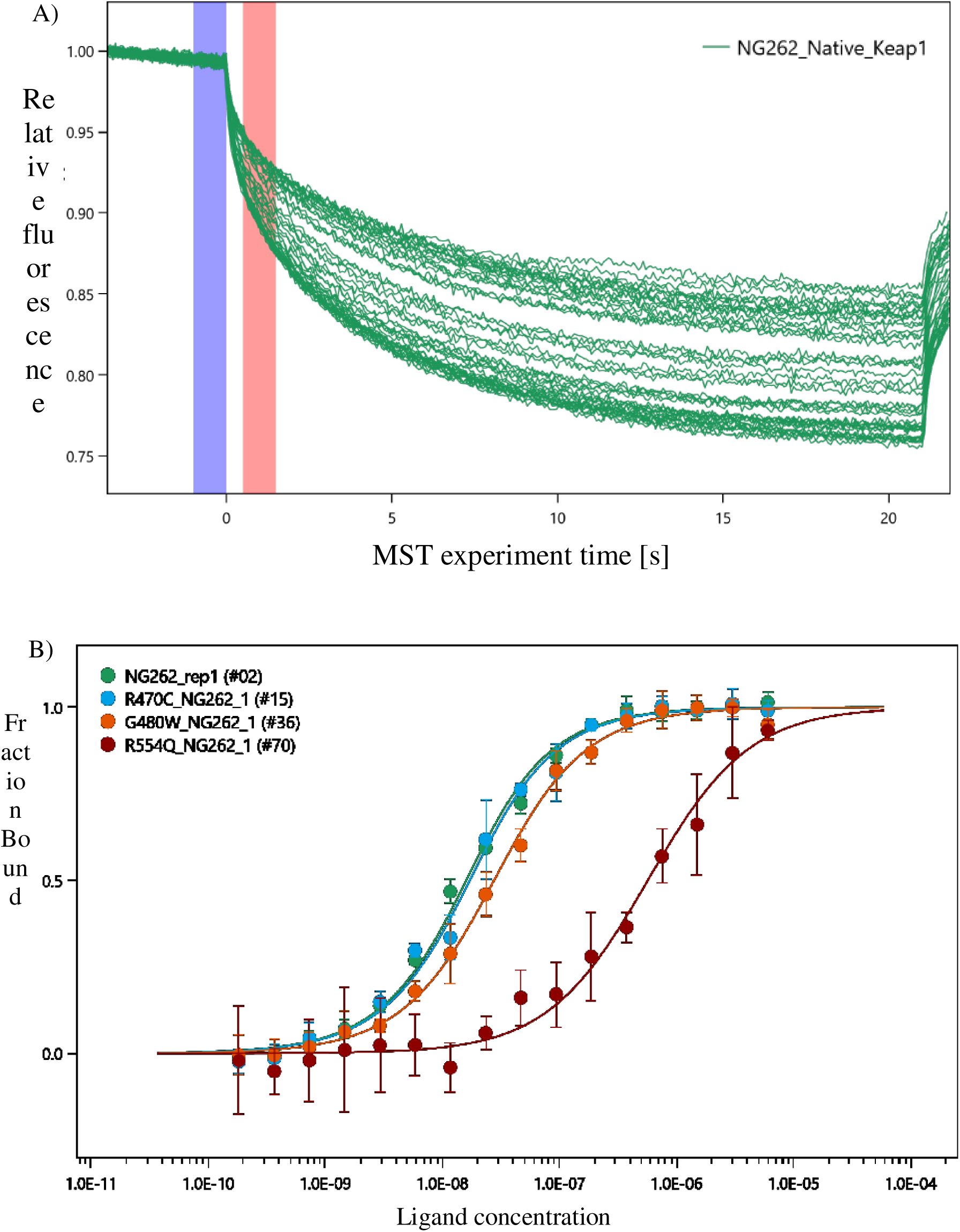
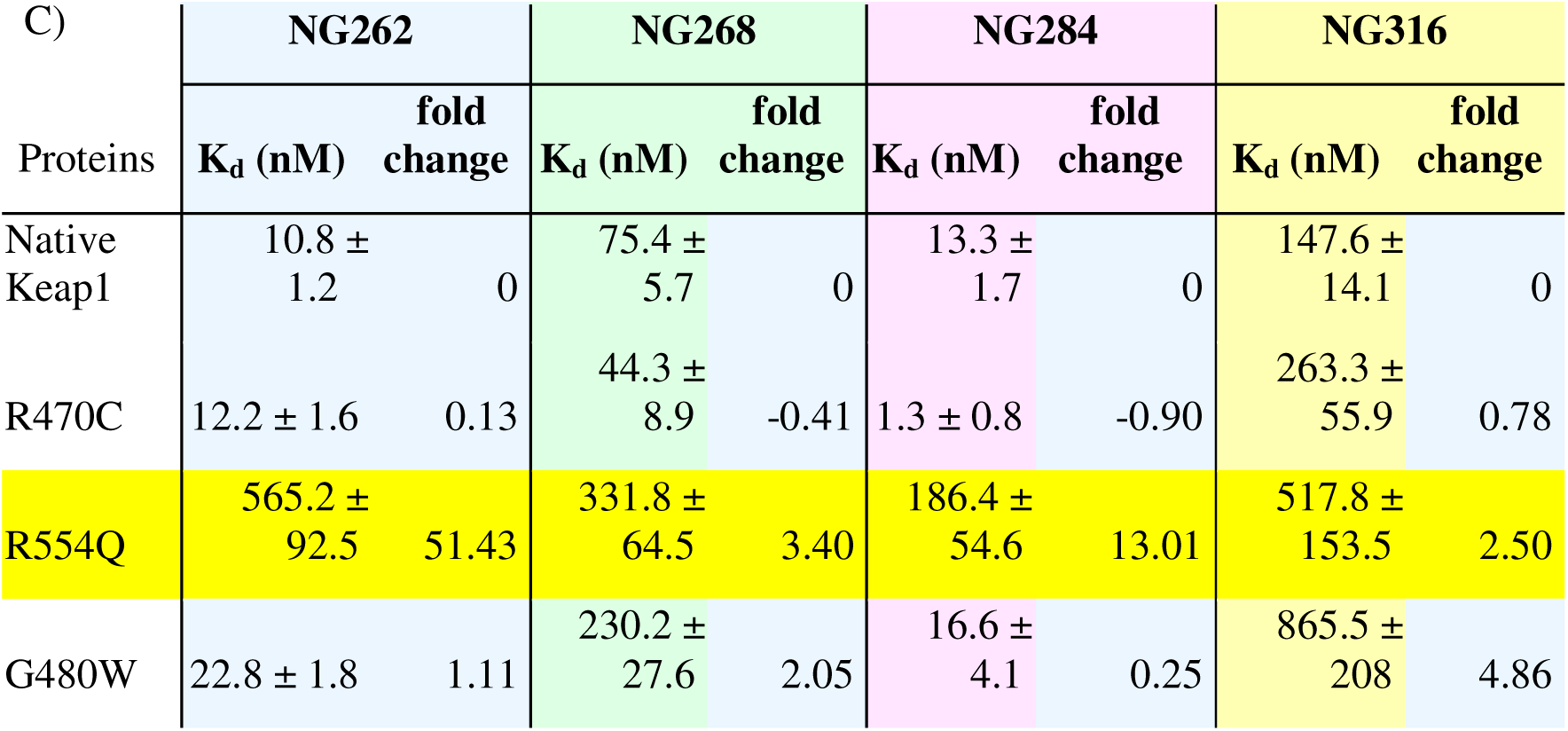
A) Exemplified MST-traces of native Keap1 with NG262; blue region represents F_0_ (cold), red region represents F_1_ (hot). B) Dose-response curves of native Keap1and its mutants with NG262. C) Binding affinities of native and mutant Keap1 for the four inhibitors.

The R470C mutant exhibited a binding affinity range comparable to native Keap1 (Fig 8C). However, the G480W mutant displayed slightly weaker binding affinities to NG262 and NG284, with a change of less than two-fold in the dissociation constant. The G480W mutant exhibited significant decreases in affinity for NG268 and NG316, with twofold and fivefold increases in K_d_, respectively. In the R554Q mutant, binding affinities for all four inhibitors were markedly reduced. R554Q-NG262 showed the highest fold change, with a K_d_ of 565.2 ± 92.5 nM, representing a 51-fold increase, followed by a 13-fold change for NG284, with a K_d_ of 186.4 ± 54.6 nM. The R554Q mutant exhibited approximately three-fold and two-fold changes for NG268 and NG316, with K_d_ values of 331.8 ± 64.5 nM and 517.8 ± 153.5 nM, respectively. In summary, MST results highlighted that the R554Q mutant had the most significantly altered binding affinities, particularly for NG262 and NG284, while G480W showed moderate changes and R470C retained affinities similar to native Keap1.The reduced inhibitor binding affinities observed for G480W and, more markedly, for R554Q are consistent with the corresponding loss of thermostability detected in TSA experiments. This suggests that decreased protein stability may contribute to the diminished ligand-binding capacity in these mutants. Consequently, such reductions in binding affinity are likely to impair the efficiency of these inhibitors in competing with NRF2 and potentially limiting their functional inhibitory activity.

### Spatial distribution of mutations suggests functional clustering in the Kelch domain

To better understand the structural and functional landscape of KEAP1 mutations, we examined their spatial distribution across the full-length KEAP1 architecture. Superbinder mutations, which exhibit enhanced NRF2 binding, are predominantly localised within or adjacent to the intervening region (IVR). Aside from G186R and S243C, located in the BTB and IVR domains, the remaining superbinder residues, R320Q, D422N, G423V, and R470C are clustered on a specific face of the KELCH domain. Although R320Q lies outside the crystallised region, upstream of the resolved KELCH domain, D422N, G423V, and R470C localise proximally on the same surface of the KELCH β-propeller, facing toward the IVR domain. This clustering may indicate an allosteric interface that facilitates domain communication, potentially modulating binding affinity through local flexibility or conformational regulation (Fig 9). In contrast, loss-of-function mutations including R554Q, G333C, W544C, and N469fs are primarily situated within the KELCH domain, particularly in regions involved in maintaining structural stability or direct interactions with NRF2. These mutations likely impair the binding interface or disrupt local folding, resulting in reduced or abolished NRF2 binding. Hypomorphic mutations, which retain partial binding capacity, are found in both the BTB domain (V167F) and the KELCH domain (G480W), suggesting that moderate functional disruption can arise from mutations in either direct or distal domains. This underscores the modular but interconnected nature of KEAP1 domain functions. To visualise the broader spatial context of these mutations, the full-length KEAP1 structure was predicted using AlphaFold3, which provided high-confidence modelling across the entire protein. The crystal structure of the KELCH domain was overlaid onto the predicted model to validate domain alignment and aid in mutation mapping. All 18 cancer-associated KEAP1 mutations reported in lung squamous cell carcinoma were then mapped onto this model and colour-coded according to their functional classification (Fig 7). This structural mapping highlights the surface clustering of superbinder mutations, the localisation of loss-of-function variants in sensitive structural regions, and the broader distribution of passenger and hypomorphic mutations. Collectively, these findings suggest that the KELCH domain is structurally resilient, with most point mutations not inducing major conformational changes. Instead, functionally relevant effects may arise from perturbations in flexible regions such as the IVR, or from altered allosteric interactions between domains. The clustering of superbinder mutations on a specific KELCH surface suggests a previously underappreciated regulatory axis for modulating KEAP1–NRF2 binding dynamics. In summary, our analysis reveals a distinct domain-specific pattern in the functional classification of mutants.

**Figure 9.**
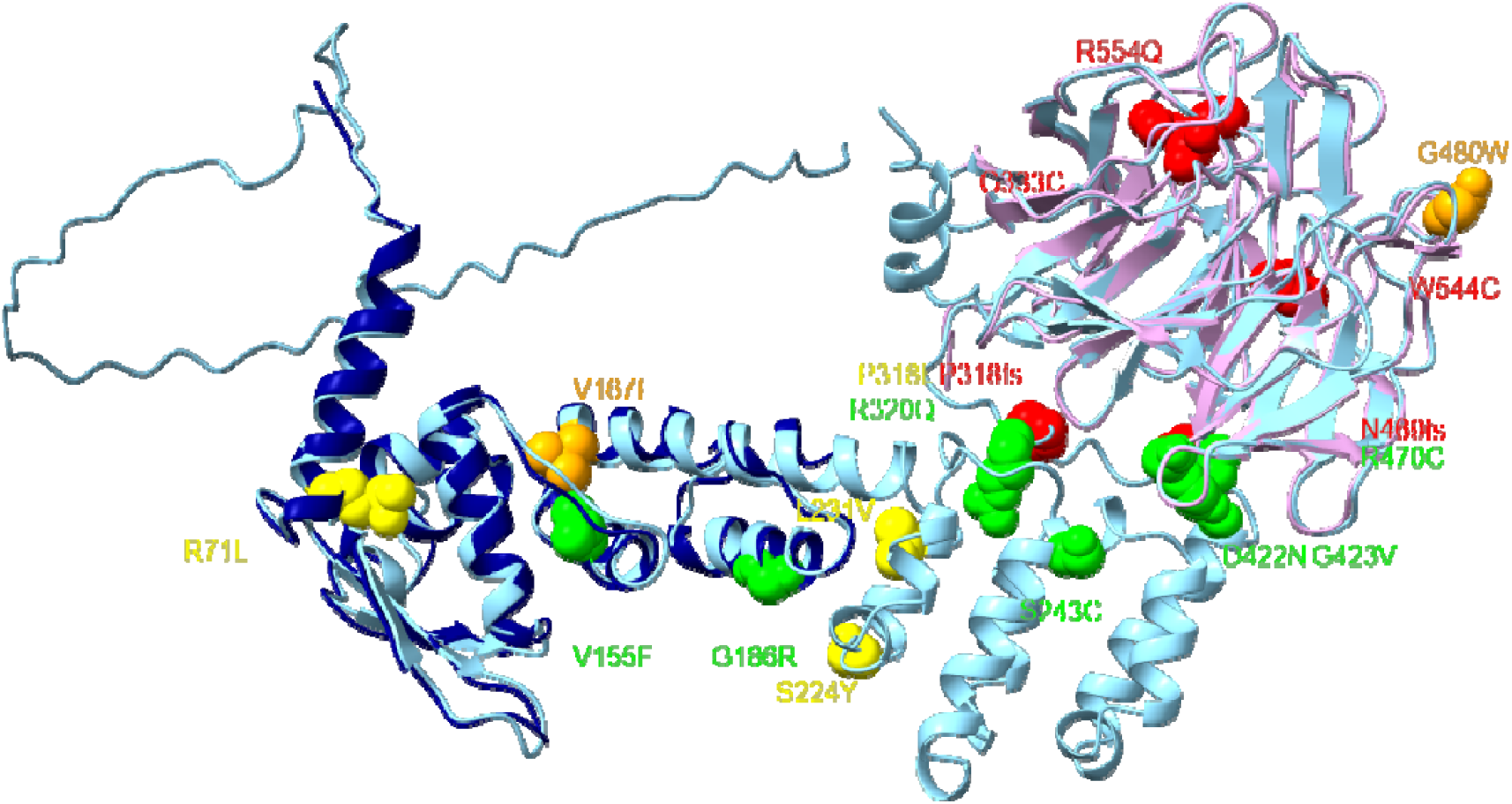
Structural mapping of cancer-associated mutations onto the predicted full-length Keap1 structure. The KEAP1 structure predicted using AlphaFold3 (light blue), and the experimentally resolved crystal structures of the BTB domain (dark blue, PDB entry 5NLB) and KELCH domain (pink) were overlaid for reference. All 18 reported KEAP1 mutations in lung squamous cell carcinoma are shown as spheres and colour-coded by functional classification: green colour indicates superbinder mutations, red colour shows loss-of-function mutations, orange colour highlights hypomorphic mutations, and yellow displays passenger mutations.

## Conclusions

This study provides an integrated structural and biophysical characterisation of three cancer-associated KEAP1 Kelch domain mutants, R470C, G480W and R554Q, with implications for their differential regulatory role in NRF2 signalling. Biophysical assays confirmed that these mutants exhibit distinct stability profiles and binding affinities for both NRF2-derived peptides and small-molecule inhibitors.

The R470C mutant, classified as a “superbinder”, retained thermal stability and showed enhanced binding affinity for the ETGE motif of NRF2. Interestingly, despite this increased binding affinity, previous reports have shown that the R470C mutant facilitates NRF2 ubiquitination but fails to promote its subsequent proteasomal degradation. As a result, NRF2 is aberrantly stabilised in the presence of R470C compared to wild-type KEAP1 or the hypomorphic G480W mutant (Hast, B.E., *et al*., 2014). This phenotype highlights a mechanistic decoupling between NRF2 binding and degradation, suggesting that R470C may impair downstream ubiquitination complex dynamics or conformational transitions required for efficient substrate processing. In contrast, R554Q, identified as a null allele exhibited a substantial loss of thermal stability and markedly reduced binding affinity, consistent with its inability to effectively interact with NRF2. The G480W mutant, a hypomorphic variant, displayed moderately decreased stability and binding affinity, representing a partial loss of function.

Crystal structures of the KEAP1-inhibitor and KEAP1-ETGE peptide complexes revealed that all three mutants adopt conformations highly similar to the native protein, with preserved inhibitor-binding orientations and peptide engagement. Despite the amino acid substitutions, no significant global conformational shifts were observed within the Kelch domain. However, local side-chain interactions were altered, particularly in the R554Q mutant, where disruption of hydrogen bonding at the C-terminal subdomain interface may underlie the pronounced destabilisation.

Spatial mapping of known cancer-associated mutations suggested functional clustering, with superbinder mutations predominantly localized near the IVR interface, while most loss-of-function mutations were found within the Kelch domain. Structural modelling using AlphaFold3 further supported these observations, indicating that conformational changes may be confined to flexible or non-crystalline regions, such as the IVR domain. Together, these findings offer valuable insight into how KEAP1 mutations influence protein stability, NRF2 binding, and drug engagement, advancing our understanding of the molecular mechanisms underlying KEAP1 dysfunction in cancer. This knowledge may inform future therapeutic strategies aimed at modulating the KEAP1-NRF2 pathway.

## Abbreviations

ARES: Antioxidant Response Elements
BTB: Bric-à-Brac
DMSO: dimethyl sulfoxide
DTT: Dithiothreitol
HEPES: 4-(2-Hydroxyethyl)piperazine-1-ethanesulfonic acid
IPTG: Isopropyl β-D-1-thiogalactopyranoside
ITC: Isothermal titration calorimetry
IVR: intervening region
Keap1: Kelch-like ECH-associated protein 1
MST: Microscale thermophoresis
NKT: natural killer T cells
NRF2: Nuclear factor erythroid 2-related factor 2
PMSF: phenylmethylsulfonylfluoride
ROS: reactive oxygen species
PTMs: post-translational modifications
PTMs: post-translational modifications
ROS: reactive oxygen species
SPR: surface plasmon resonance
SQCC: squamous cell lung carcinomas
TCGA: Cancer Genome Atlas
TSA: Thermal shift assays

## Acknowledgements

This project was supported by the National Science and Technology Development Agency (NSTDA), Thailand, through the Royal Thai Government scholarship programme. G.W. would like to acknowledge financial support from Cancer Research UK (C9344/A10268), the Bloomsbury Consortium, and UCL School of Pharmacy.We also acknowledge Diamond Light Source (Didcot, UK) for access to macromolecular crystallography beamlines under proposals mx38316-13 and mx30393-53, which enabled data collection for this study. This work contains part of the doctoral theses of Qian Wang and Watanyoo Sopipong.

